# Effect of Endoplasmic Reticulum Stress on Human Trophoblast Cells: Survival Triggering or Catastrophe Resulting in Death

**DOI:** 10.1101/2021.09.15.460465

**Authors:** Gurur Garip, Berrin Ozdil, Duygu Calik-Kocaturk, Fatih Oltulu, Fatma Zuhal Eroglu, Huseyin Aktug, Aysegul Uysal

## Abstract

Although *in vitro* endoplasmic reticulum (ER) stress studies have been carried out using Tunicamycin in human trophoblast cell lines in recent years, the effect of calcium homeostasis impaired by the effect of Thapsigargin on cell survival - death pathways have not been clearly demonstrated.

Here, the effects of ER stress and impaired calcium homeostasis on cell death pathways such as apoptosis and autophagy in 2-dimensional and 3-dimensional cell cultures were investigated using the HTR8 / SVneo cell line representing human trophoectoderm cells and the ER stressor Thapsigargin. By using Real Time PCR, gene and immunofluorescence analyzes were studied at the protein level.

In this study, it has been established that the Thapsigargin creates ER stress by increasing the level of GRP78 gene and protein in 2 and 3 dimensions of human trophoectoderm cells and that cells show different characterization properties in 2 and 3 dimensions. It has been determined that while it moves in the direction of EIF2A and IRE1A mechanisms in 2 dimensions, it proceeds in the direction of EIF2A and ATF6 mechanisms in 3 dimensions and creates different responses in survival and programmed cell death mechanisms such as apoptosis and autophagy.

With forthcoming studies, it is thought that the effects of Thapsigargin on the intrinsic pathway of apoptosis and the linkage of the autophagy mechanism, the examination of the survival-death pathways in the co-culture model with endometrial cells, therapeutic target molecules that will contribute to the elucidation of intracellular cell dynamics may increase the success of implantation.

## 1. INTRODUCTION

World Health Organization (WHO) defines infertility as “the inability to achieve clinical pregnancy despite regular unprotected sexual intercourse for 12 months or more”. Infertility is a major health problem worldwide, estimated to affect 186 million people worldwide (Elhussein et al., 2019). Implantation failure with unknown factors takes a large place in infertility.

Preimplantation period constitutes the first stages of the development of mammalian embryos in the uterus. In preimplantation period, early blastomeres divide and differentiate. The first differentiation of the embryo before implantation causes the blastocyst which is consisted of inner cell mass (ICM) and trophectoderm (TE) to emerge. Trophoectoderm cells consist of the epithelial layer that provides the first contact with the uterine wall (De Paepe et al., 2018). This cell layer contains trophoblast stem cells, cytotrophoblast, syncytiotrophoblast, extravillous trophoblast cells, cell arm, endoglandular and endovascular trophoblast cells (Gamage et al., 2016). Placenta differentiates from the trophoblastic cell line that will contribute to the extraembryonic part of the chorion. Disruption in the specification of the trophoectoderm leads to impaired embryo implantation and constitutes the leading cause of infertility (De Paepe et al., 2018).

Cytotrophoblast, syncytosisotrophoblast cells and trophoblastic stem cells is represented by the HTR8/SVNeo cell line originating from first trimester extravillous healthy trophoblast cells whose division ability was enhanced by large T antigen transfection Simian Virus 40, an oncogenic protein. Studies have revealed that this cell line contains a heterogeneous cell population. When the cell line is examined by flow cytometry, CK7 or E-cadherin positive epithelioid trophoblast cells, HLA-G positive extravillous trophoblast cells and Vimentin positive mesenchymal trophoblast stem cells can be detected (Abou-Kheir et al., 2017) (Msheik et al., 2020). It was determined cells that contain trophoblast cells (CK7+ and CSH+) containing extravillous trophoblast markers HLA-G and ITGAV/ITGB3, villous syncyrotrophoblast marker CGB, PHLDA2 and villous trophoblast progenitor marker TCL-1. The cell line staines positive with Hoechst 33342 and expresses ABCG2 and ID2. These markers show that there are progenitor stem cells in the cell line. It has been shown that progenitor cells express BMP4, which is an important stage in differentiation from embryonic stem cell to trophoblast stem cell (Takao et al., 2011). In another study, it was determined that the cell line expressed genes with stem cellular characteristics such as SOX2 and NANOG. It has been attested that the trophoblast stem cell marker CDX2 and the human embryonic stem cell marker NOTCH1 are positive (Weber et al., 2013). In studies conducted, when the cell line was examined and the receptors of the cells were evaluated, the model exactly reflects the reality in imitating the implantation stage of a healthy human embryo (Hannan et al., 2010). Construction and destruction processes continue throughout the life cycle of cells. Proteins synthesized from ER ribosomes are transformed into mature proteins by undergoing some post-translational modifications in the ER (Adams et al., 2019). During the cell life process, there may be misfolded protein accumulation in the ER. Misfolded proteins in the ER are directed to correct folding with the Unfolded Protein Response (UPR), primarily through GRP78, GRP94, calnexin, calreticulin and calcium-dependent chaperones. Since protein folding errors cannot be corrected, the amount of misfolded protein in the ER lumen increases and causes ER stress. When ER stress occurs, the cell tries to correct the misfolded proteins that cause stress by reducing the transcription of the misfolded proteins and inducing autophagy with the help of chaperones. Activated chaperones prompt the ways to relieve stress by activating different mechanisms and signal pathways. Incorrectly folded protein accumulation, which cannot be corrected even with the activation of all these pathways, can direct the cell to apoptosis (Iurlaro et al., 2016).

It has been indicated that the ER stress that occurs in the cell plays a role in the pathogenesis of many diseases such as type 2 diabetes, Parkinson’s, retinitis pigmentosa, Alzheimer’s, and multiple myeloma (Ozcan & Tabas, 2012). Studies on the pathogenesis of ER stress in implantation biology are limited and its role has not yet been established. It has been studied ER stress-induced responses play critical role for embryo development. It has been shown that increased ER stress in decidual tissue in pregnancy has been shown to be implicated in fetal growth restriction with and without pre-eclampsia. Sustained ER stress acts as a cofactor of oxidative stress in decidual cells from patients with early pregnancy loss (Michalak & Gye, 2015). When the literature is examined on human trophoclastic cell line; propyl gallate, used on the HTR-8/SVneo line which is an additive in the food and cosmetic industry, affects proliferation, apoptosis, and invasion behaviors (Yang et al., 2017). Regarding the effects of butyl paraben, which is frequently used in shampoo, cream and make-up materials, it has been reported that it affects proliferation, apoptosis, invasion behaviors by increasing ROS on the HTR-8/SVneo line in placental development (Yang et al., 2018). In the review written by Güzel et al, the effects of ER stress on the urogenital system were explained and it was stated that more studies should be done (Guzel et al., 2017). Thapsigargin, which is used as an ER stressor, has been studied in many cell lines such as neuronal stem cells, melanoma, hematopoietic, pancreatic beta, and prostate cancer. Studies on the effects of Thapsigargin on the urogenital system involving the HTR-8/SVneo cell line are limited. Based on these data, we determined the question of our study as what the effect of ER stress is created on trophectoderm cells on intracellular traffic and whether this emerging problem can be solved with therapeutic targets. The hypothesis of this study is that human trophectoderm cells, one of the components of implantation, are related to changes in intracellular dynamics due to ER stress; It affects cell survival mechanisms such as apoptosis and autophagy. Here, we induced ER stress with Thapsigargin and investigated changes in the apoptosis and autophagy pathways at RNA and protein levels. This study gives clue of ER stress and calcium homoestasis importance in implantation success.

## 2. MATERIAL AND METHODS

### 2.1 Study Design

To show the behavior of cells and interactions between cells, we designed two control and two experimental groups. In the first control group, cells were cultured as 2-dimensional in a petri dish, while in the first experimental group which were also cultured as 2-dimensional in a petri dish. The effects of ER stress created through ER calcium channels by applying ER stressor (Thapsigargin) to these cells on intracellular traffic, changes in cell survival mechanisms and behavioral differences because of communication between cells were investigated. In the second control group, cells were cultured as spheroids to mimic the structure of the trophoectoderm layer in the human embryo (an average of 2000 cells per spheroid to ensure dimensional similarity). The behavior of the spheroid cells and the interaction between cells were examined and the differences in 2 and 3 dimensions were demonstrated within the framework of the panels indicated. In the second experimental group, ER stressor (Thapsigargin) was applied to 3-dimensional spheroidized cells to examine the effects of ER stress on human trophoectoderm layer mimicked in vitro. Cellular and cell-to-cell interactions that will occur with the impairment of calcium homeostasis via ER calcium channels as a result of ER stress were examined.

### 2.2 Cell Culture

#### 2.2.1 2D Cell Culture

Human trophoectoderm cells were represented by HTR-8 / SVneo cell line (ATCC CRL-3271) which is immortalized by Simian Virus 40 (SV40) viral DNA transfer of first trimester villous explants. In experiments, HTR8 / SVneo cell line cells were grown in RPMI-1640 medium (ATCC 30-2001) containing 10% Fetal Bovine Serum (ATCC 30-2020), 1% Penicillin-Streptomycin (Pen-Strep) (Gibco 15140148). Passage 4 cells were used in all experiments. Cultured cells were kept at 37°C, 5% CO2 (Abou-Kheir et al., 2017).

#### 2.2.2. Spheroid Formation with Hanging Drops

To mimic 3D human trophectoderm layer, cells were applied spheroid formation hanging drop method. Briefly, 2×10^5^ cells per 20 *μ*L drop supplemented RPMI-1640 with %10 FBS and %1 Pen-Strep were plated onto the lid of Petri dishes in regular arrays (20 drops/Petri lid). The lid was inverted over the bottom of the PBS-filled Petri dish (Weber et al., 2013) (Figure 1). The petri dish with the hanging drops were cultured under standard conditions (37°C, 5% CO_2_, humidified atmosphere) for 240 hours to imitate human 5th day blastocyst size (170-220 µm). Spheroids were washed gently with 150 mL RPMI-1640 with %10 FBS medium and transferred to petri dish coated with agarose gel (Himedia GRM026). Changes in spheroids caused by Thapsigargin exposure were assessed by RT-PCR and immunofluorescent staining methods (Winship et al., 2018).

**Figure 1:**
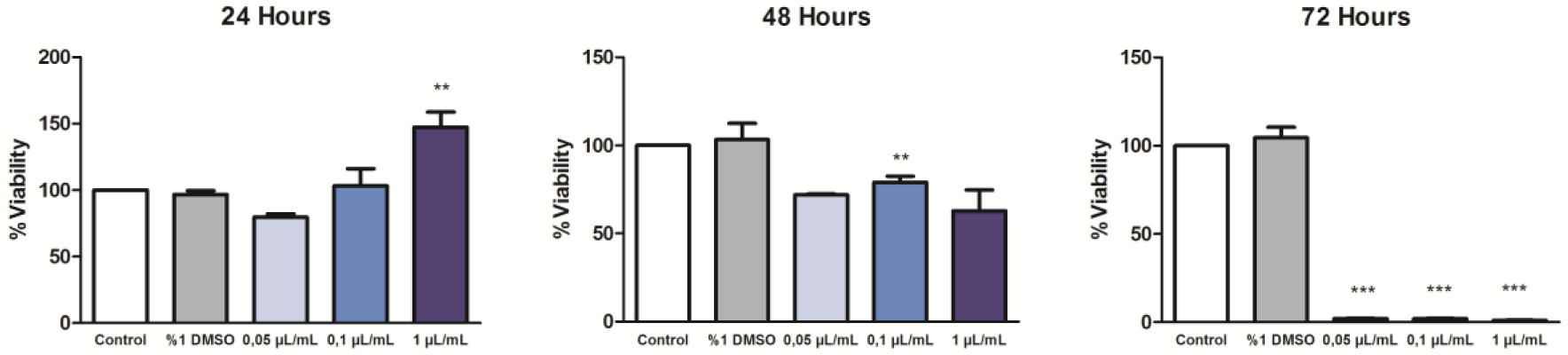
24, 48 and 72 hours Thapsigargin MTT viability results of HTR8/SVneo cells (**:p<0,05 and ***:p<0,01 Oneway ANOVA – Bonferoni post hoc test). To induce ER stress on HTR-8/SVneo cells, it was found 0.1 µL/mL Thapsigargin for 48 hours is effective.

### 2.3. Determination of Thapsigargin Effective Dose (EC50)

To determine the most appropriate concentrations of Thapsigargin (Invitrogen Cat. No # T7458), MTT Cell Proliferation Assay Kit (Colorimetric) (Biovision Catalog # K299-1000) and The Muse**®** Count & Viability Kit (Luminex Part Number: MCH100102) was used according to the manufacturer’s protocols.

#### 2.3.1. MTT Colorimetric Cell Proliferation Assay

HTR8/SVneo cells prepared as passage 4 in 96-well plate were cultured as 2×10^5^ per mL and left to incubate for 24 hrs. Thapsigargin solution was prepared at concentrations of 1-0,1-0,05 µL/mL and added to wells with equal number of cells. To show the internal control (DMSO) effect on the cell, the same amount (10 μL) of DMSO was mixed with RPMI-1640 (990 μL) medium and was added to the DMSO wells. Separately, the medium was added to a separate well as a control group. All groups were incubated for 24 hrs before measurement. Primarily, all of the mediums were removed from the tube wells and then the ‘blank’ well was added to these wells as a background control. Later on, 50 μL of MTT reagent (Biovision lot: 3K12K02991) and 50 μL of serum-free medium mixture were added to these wells and left to incubate for 3 hours at 37°C. At the end of 3 hours, 150 μL of MTT Solvent (Biovision lot: 3K12K02992) was added to all wells and placed in the mixer for 15 min. The results were obtained by studying at 590 nm in the Ege University Medical Biology Laboratory in a dark environment. The obtained MTT values were calculated by regarding the blank value. Accordingly, the percentage of other cells was calculated so that the control group was 100%.

#### 2.3.2. Muse^®^ Cell Viability Assay

HTR8/SVneo cells were cultured in 6-well plates at a density of 2×10^5^ per milliliter. 0.05-0.1-1-5 µL/mL doses of Thapsigargin were added to the cells that left in incubation for 24 hours and the cells were incubated for another 24, 48 and 72 hours. After this step, the normal passage protocol, in which dead and living cells were taken together, was applied to the cells. The obtained cells were compared using the cell number and Muse^®^ Count & Viability Kit in the Muse^®^ analyzer.

### 2.4 RNA extraction, reverse transcriptase, and quantitative real-time RT-PCR

Each group in the experimental setup was repeated in 3 times and gene analysis were carried out. The genes HSPA5, ATF6, ERN1, EIF2A from the ER stress panel (Yand et al., 2018); BCL2, CASP3 from the apoptosis panel (Gu et al., 2018); MAPLC3B, BECN1 from the autophagy panel(Q. Zhang et al., 2017) were examined in the control and experimental groups determined. The cells were removed from the flask, dissolved in RNA buffer, and stored at - 80°C. RNA samples were isolated with an RNA isolation kit (Roche Product No: 11828665001). Purity and concentration measurements were taken with nanodrop at 260-280 nanometers. RNA concentrations were diluted appropriately, and cDNA synthesis was performed with the cDNA synthesis kit (Roche Product No: 11483188001). ΔCt values were measured using the “Multiple Plate Analysis” program. The expressions of the reference genes (GAPDH, RPLPO, TBP) were used for normalization and change comparisons between groups were made using Student t-test.

### 2.5 Immunofluorescence staining and fluorescence microscopy

#### 2.5.1. Antibodies and reagents

Anti-EIF2A (BS-3613R), anti-IRE1a (BS-8680R), anti-c-Caspase3 (Bioss BS-0081R), anti-Beclin1 (S-1353R) primary antibodies were prepared within 1% BSA at 1/200 dilution, while secondary antibody Bioss Alexa Fluor-488 was prepared at 1/400 dilution manufacturer’s instructions.

#### 2.5.2. Immunofluorescence staining

##### 2.5.2.1 2D Immunofluorescence Staining

The cells were fixed in 4% paraformaldehyde for 30 min at room temperature and washed three times with 1X PBS. To increase the membrane permeability, the cells were kept in a fresh-prepared 0.25% Triton-X100 solution for 15 min and rinsed with 1X PBS. Nonspecific reactivity was blocked by incubating the cells in blocking solution (1% BSA in 1X PBS) for 1 hr in room temperature. Subsequently, the cells were subjected to primary antibody (1/200 diluated with %1 BSA) treatment for overnight at 4°C. The following day, the cells were washed 3 times with 1X PBS to remove the primary antibody. Secondary antibody (Alexa Fluor-488) was prepared at 1/400 dilution as specified by the manufacturer. Cells were washed with 1X PBS by soaking in secondary antibody for 1 hr. After washing with 1X PBS, slides were mounted with a drop of mounting medium (with Fluoroshield Mounting Medium with DAPI (Abcam, ab104139)) and images were captured under Olympus BX50 fluorescence microscope.

##### 2.5.2.2 3D Immunofluorescence Staining

The control and experimental group spheroids were applied by using the solutions included in the 2D immunofluorescence staining protocol, and the spheroids were transferred to the wells containing the solutions and kept for the waiting period specified in the protocol (Balahmar et al., 2018).

### 2.6. Statistical analysis

One-way ANOVA and post hoc Bonferonni tests were performed for MTT data. The expressions of the reference genes (GAPDH, RPLPO, TBP) were used for normalization and change comparisons between groups were made using Student t-test. At least 100 cells were taken into considiration for protein intensity measurements and the values ‘integrated intensity’ were obtained by ImageJ. These values were evaluated by GraphPad Prism 8.4.3 analysis software statistically and analyzed with Student t-test. The average protein intensity rates obtained from the x and y axes of the spheroids were compared with the control groups and the radiation intensities were displayed graphically in the same way of 2-dimensional study. All experiments were replicated three times.

## 3. RESULTS

### 3.1. Effective Dose of Thapsigargin to ER stress in HTR-8/SVneo cells

As a result of MTT experiments, it was found that 0.05, 0.1 and 1 µL/mL doses of Thapsigargin did not have a significant effect on cell viability at 24 hours of exposure, reduced cell viability to 70% at 48 hours and decreased cell viability to 1% at 72 hours (Figure 1).

Analysis of viability and cell counts of Thapsigargin doses at 24, 48 and 72 hours were performed with the Muse^®^ Cell Viability Kit. This analysis verified the results of MTT results. Drug resistance was found in the 72-hour group. As a result, the effective dose of Thapsigargin was 0.1 µL/mL and should be administered for 48 hours for ER stress induction on HTR-8 / SVneo cell line (Figure 1).

### 3.2. Human Trophoectoderm Layer Mimicry

The experimental setup was carried on 50 spheroids from the control group and the experimental group. At the end of the 240 hours, it was determined that the density of some spheroids in the experimental group decreased, their shape was distorted, and the cells separated in their surroundings. RT-PCR and immunofluorescence experiments were continued by selecting 42 pieces of spheroid assemblages whose shape was not deformed, with ideal formation and maturation.

Spheroid structures were evaluated by morphological structure on agar and their diameters by examining with Phase Contrast Microscope (Olympus CX31RTSF-5). It was observed that the spheroids in the control group had a small structure and were denser and smaller than the experimental group spheroids. At the same time, it was observed that in the proliferation zones of the experimental group spheroids some of the cells were poured into the agar and separated from the spheroid, also there were irregularities and protrusions in their shapes (Figure 2).

**Figure 2:**
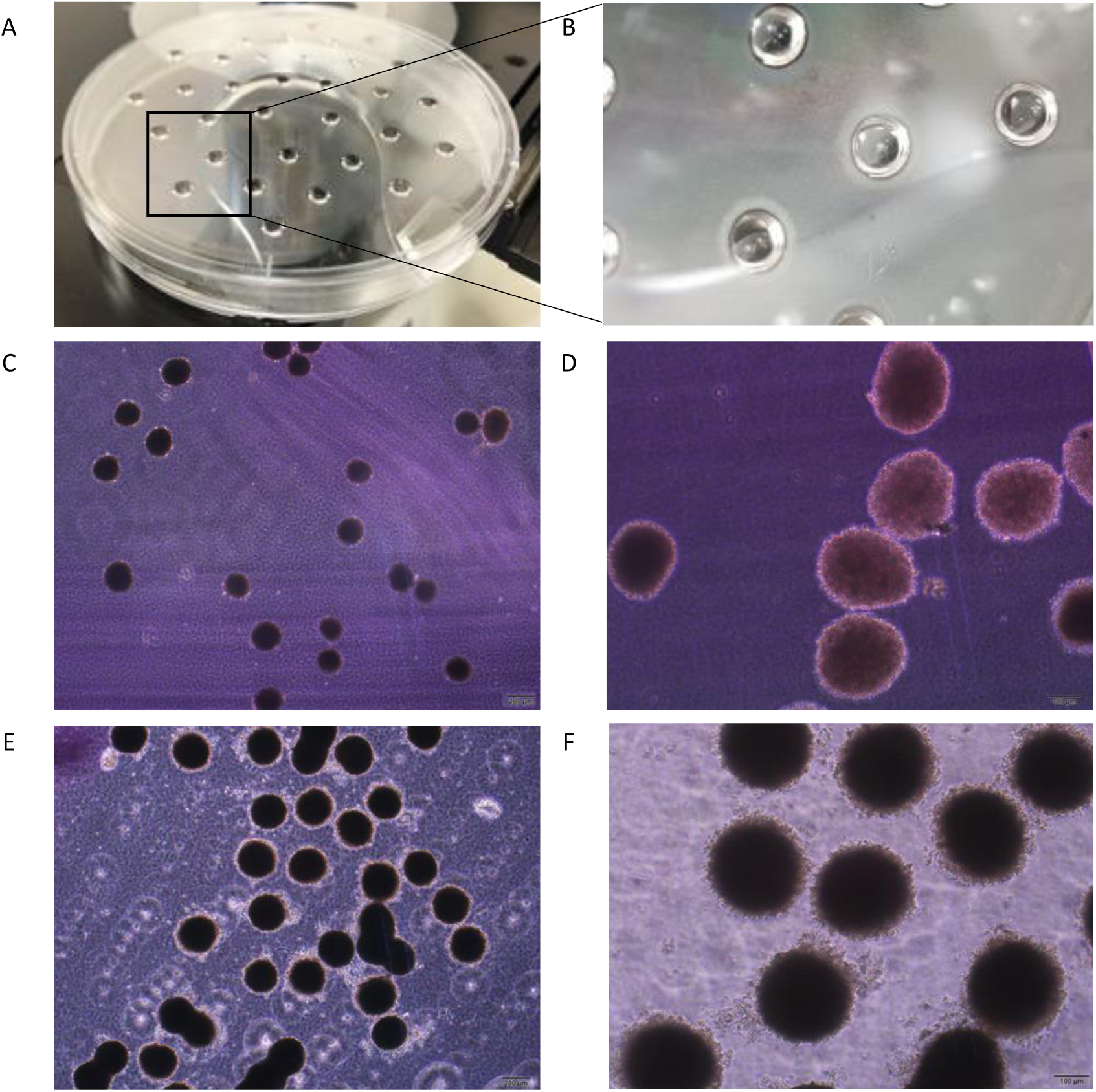
A) and B) The hanging drop model in spheroid cell culture. C) and D) Phase contrast microscope images of control group spheroids on Agar. Scale bar 100µm. E) and F) Phase contrast microscope images of experimental group spheroids on agar. Scale bar 100µm.

### 3.3. Gene Expression Analysis Findings

As a result of gene expression analysis, “Average ΔCt” rates were obtained by subtracting the reference gene rates from the rates obtained for the genes investigated for the 2- and 3-dimensional experimental group. In order to determine the increase, decrease or stay the same in the rates obtained with these results, the results of “fold regulation” were obtained by comparing the numerical data with the 2- and 3-dimensional control groups and the rates with a minimum 2-fold change in the positive or negative direction were considered significant (Figure 3).

**Figure 3:**
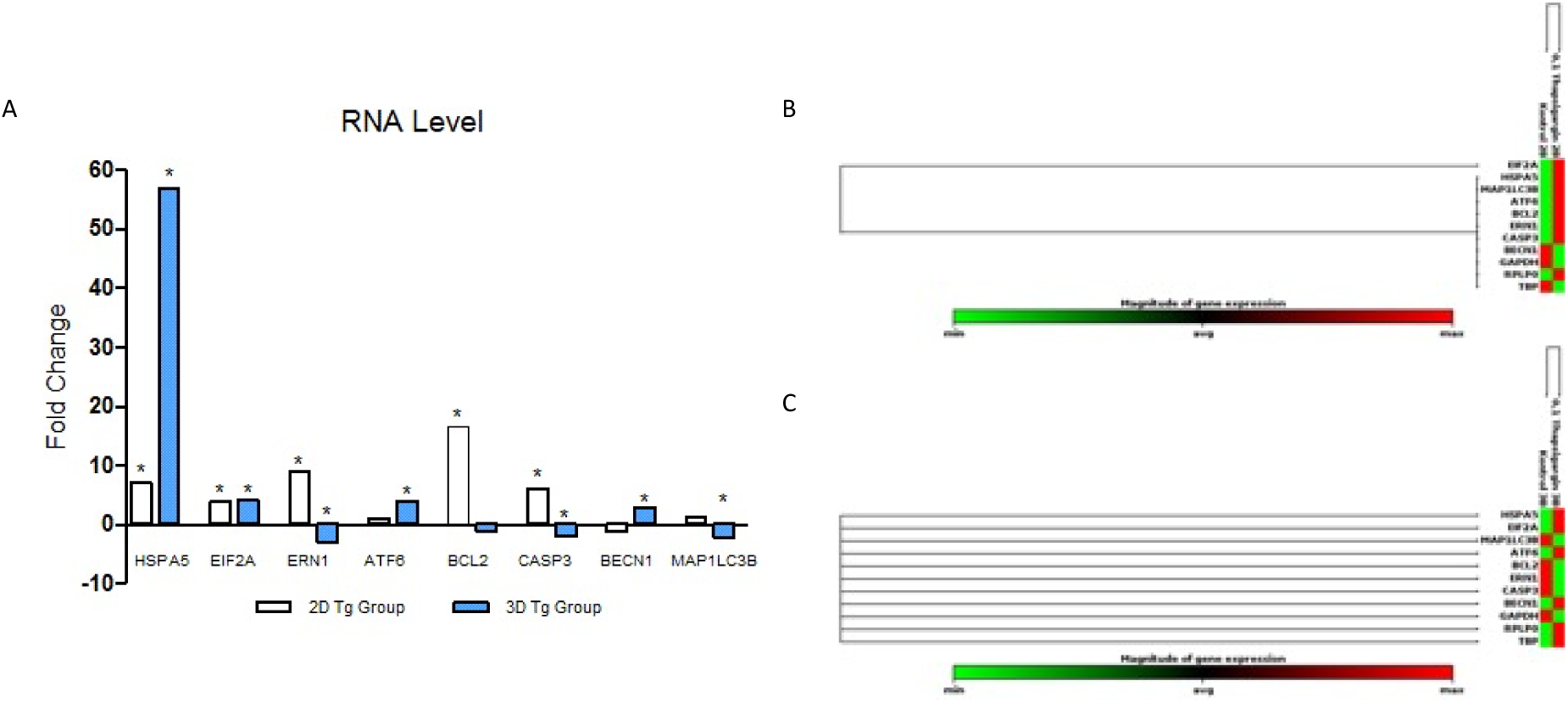
A) RT-PCR showing the expression of ER stress, apoptosis and autophagy genes in 2D and 3D HTR-8/SVneo cell cultures (*:p<0,05). B) Cluster gram chart of 2-dimensional control and drug groups gene expression levels. C) Cluster gram chart of 3-dimensional control and drug groups gene expression levels

The experiments were evaluated by repeating the expression of genes HSPA5, ATF6, ERN1, EIF2A, BCL2, CASP3, MAP1LC3B, BECN1 genes associated with ER stress pathway, apoptosis and autophagy. HSPA5 gene expression showed a 7-fold upregulation in the 2-dimensional experimental group compared to the control group, and a 56.97-fold increase in the 3-dimensional spheroid experimental group. There was no significant upregulation of ATF6 gene expression in the 2-dimensional experimental group, while a 4.04-fold upregulation of ATF6 expression was observed in the 3-dimensional experimental group. It was observed that ERN1 gene expression showed 8.9 times upregulation in the 2-dimensional experimental group compared to the control group and a 2.92-fold downregulation occurred in the 3-dimensional experimental group. EIF2A gene expression was 3.83-fold upregulated in the 2-dimensional experimental group and 4.09-fold upregulated in the 3-dimensional experimental group. When the BCL2 gene expression was examined, 16.65 times upregulation was observed in the 2-dimensional experimental group compared to the control group and no significant change was found in the 3-dimensional experimental group compared to the control group. In the CASP3 gene expression, 6.14 times upregulation occurred in the 2-dimensional experimental group compared to the control group, and 2.05-fold downregulation in the 3-dimensional spheroid experimental group compared to the control group. There was no significant change in MAPLC3B gene expression between the 2-dimensional experimental group and the control group. When the 2-dimensional experimental group and the control group were compared, there was no significant change in BECN1 gene expression. In the 3-dimensional experimental group, 2.79 times upregulation was measured compared to the control group (Figure 3).

### 3.4. Immunofluorescent Staining Findings of Proteins Associated with Active Genes

Regarding the ER stress signaling pathway, in immunofluorescence staining using GRP78, IRE1α, EIF2α antibodies, protein integrated density was found to be higher in the 2 and 3 dimensional experimental groups compared to the control groups. In the immunofluorescence staining performed using c-Caspase3 antibody related to the apoptosis pathway, an increase was found in the integrated density of the 2 and 3 dimensional experimental groups compared to the control groups. In the immunofluorescence staining using the autophagy associated Beclin1 antibody, an increase in the integrated density of the 2 and 3 dimensional experimental groups compared to the controls was found (Figure 4).

**Figure 4:**
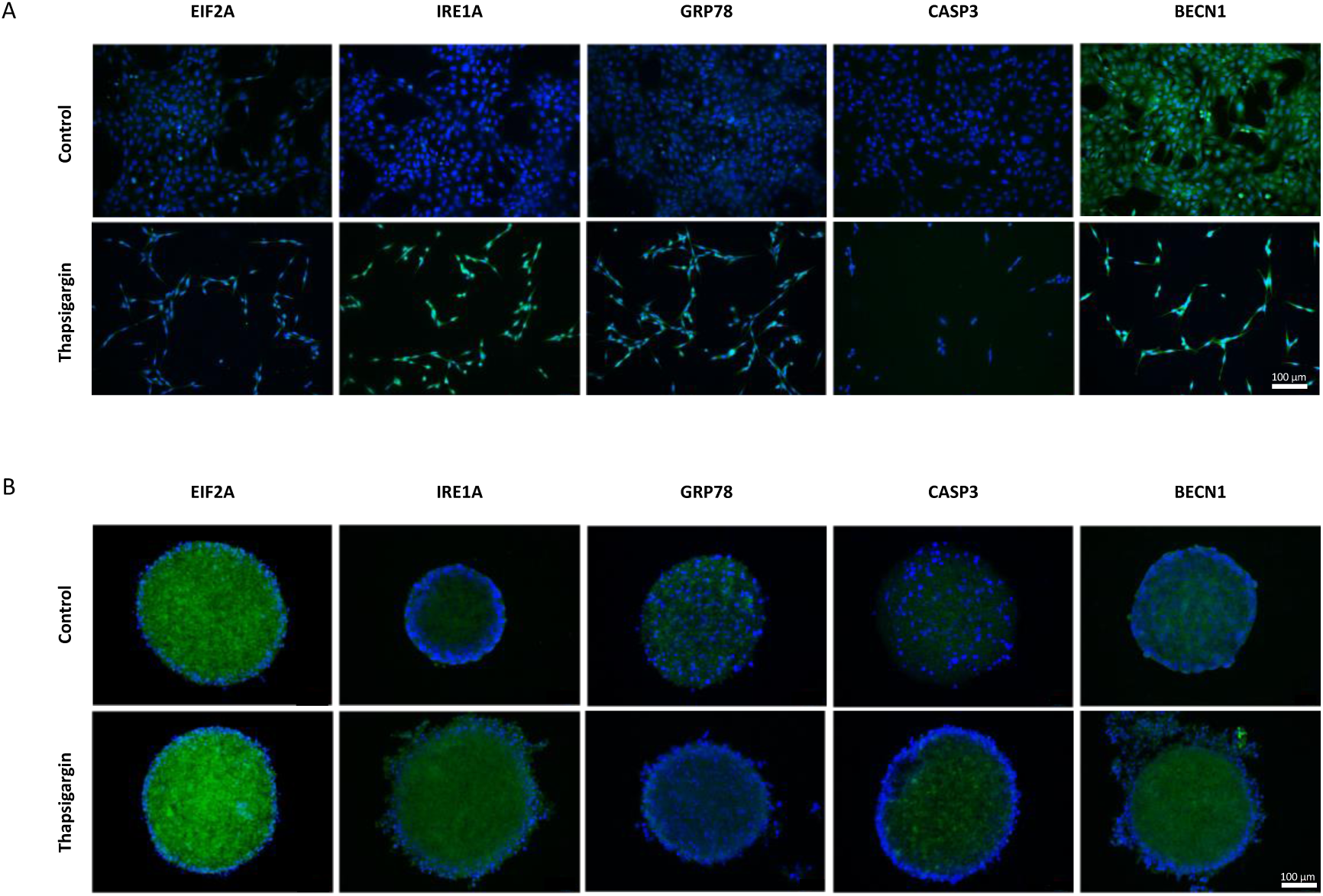
A) 2D immunofluorescent staining. Protein of interest and nucleus represented in green and blue colour in merged images, respectively. ER stress, apoptosis and autophagy signaling protein integrated density was found higher in 2D experimental groups (p<0,01). Scale bar 100µm. B) 3D immunofluorescent staining. Protein of interest and nucleus represented in green and blue colour in merged images, respectively. ER stress, apoptosis and autophagy signaling protein integrated density was found higher in 3D experimental groups Scale bar 100µm.

## 4. DISCUSSION

Implantation involves the process of a free active blastocyst attaching to the endometrium, invasion of the stroma and forming the placenta. The implantation process covers a complex sequence of events occurring in a well-defined period in which both embryo and endometrial development are synchronized. There are three prerequisites for successful implantation. These prerequisites contain an embryo capable of implantation, an endometrium in the receptive state and synchronized development between the embryo and the endometrium (S.-M. Kim ve Kim, 2017).

Although today’s scientific technology is advanced, the molecular interactions in the implantation view, implantation disorders and the molecular cellular dynamics of placental diseases have not been fully elucidated. The unethical nature of experimentation with human blastocysts and the inability to fully imitate the implantation process in vitro with receptive uterine tissue underlies these limitations. Traditional two-dimensional cell culture is limited to representing the biology of cells in vivo, and the 3-dimensional spheroid culture technique reflects cell biology and behavior more specifically. For this reason, the spheroid model, which is one of the methods used for in vitro imitation of human trophoectoderm cells in cell culture studies, is currently used in studies. (Nandi, Lim, Torres-Garcia, ve Lala, 2018) (He ve ark., 2019).

There is an increase in research on the pathophysiological roles of ER stress in various diseases. In relation to ER stress, metabolic diseases have been classified as neuro degenerative diseases, immune disorders, cancer and new ways have been opened for therapeutic approaches (Tatar ve Tatar, 2018). Yet, studies on the effect of ER stress on implantation disorder are limited. Initially, in *in vitro* studies, ER stress was created using Tunicamycin and it was studied over 2-dimensional choriocarcinoma lines. Recently, there are studies showing DNA damage in human preimplantation embryos as a result of ER stress (Dicks ve ark., 2017) and associating ER stress formation and viability as a result of intermittent hypoxia in HTR-8/SVneo cells (W. Song ve ark., 2020). Thapsigargin boost cytosolic calcium concentration by blocking the cell’s ability to pump sarcoplasmic and ER calcium. Depletion of storage secondarily activates plasma membrane calcium channels, allowing calcium influx into the cytosol. Depletion of ER calcium stores leads to ER stress and activation of unfolded protein response (Malhotra ve Kaufman, 2007).

In this study, genes associated with the ER stress pathway were evaluated with expressions of HSPA5, ATF6, ERN1 and EIF2A, and proteins associated with active genes with GRP78 (HSPA5), EIF2α, IRE1α (ERN1) immunofluorescence staining. In studies, when ER stress occurs in the reproductive system, GRP78 expression has a key role in chaperone function and UPR response formation. It has been reported that there is an increase in HSPA5 levels in pregnant women diagnosed with preeclampsia and as a result, ER stress occurring in trophoblast cells causes disruption in lysosome structures and autophagy is induced (Nakashima, Cheng, Kusabiraki, Motomura, ve Aoki, 2019). It has been demonstrated that by applying Tunicamycin and Thapsigargin to JEG-3 and HTR8 / SVneo trophoblast cells, HSPA5 expression increased in both cell lines and decreased the invasion by suppressing MMP2 mRNA and protein expression in the cells (C.-L. Lee ve ark., 2019). Steroidogenesis mechanism is arranged proliferation and cell survival by affecting the expression of GRP78 (Hebert-schuster ve ark., 2018). ATF6, which functions as an ER stress sensor, creates the cis-acting ER stress response element (ERSE) found in the promoters of genes encoding ER chaperones when ER stress occurs (Hillary ve Fitzgerald, 2018). In preeclamptic pregnant women, sFlt-1 levels change due to ER stress with activation of ATF6 mechanism of trophoblasts in placenta (Mochan, Dhingra, Varghese, ve Gupta, 2017). It has been demonstrated that ER stress in the placental trophoblast cells of preeclamptic pregnant women occurs through the ATF6 signaling mechanism, and these changes are induced by apoptosis by disruption in the nitric oxide signal pathway (Du ve ark., 2017). In the signaling processes of trophoblasts, there is a connection with autophagy through Beclin-1 expression and ER stress through the ATF-6 mechanism (Bastida-ruiz ve ark., 2019).

IRE1α protein functions as a sensor of unfolded proteins in the endoplasmic reticulum, triggering the unfolded protein response (UPR) signaling pathway (Y. Chen ve Brandizzi, 2014). Ire1α is actively found in trophoblasts, especially in the placenta, and loss of its expression leads to a decrease in VEGF-A levels, causing to implantation disorders (Iwawaki, Akai, Yamanaka, ve Kohno, 2009). Similarly, it has been shown that in Ire1α knockdown trophoblast cells, placental disorder occurs due to a significant decrease in VEGF and implantation disorder occurs with the activation of other ER stress pathways (Ghosh ve ark., 2010). In HUVEC cells under hypoxia conditions, a negative correlation was determined between Hif1-α and IRE1-α activity as an ER stress pathway (Collawn ve Bartoszewski, 2020).

EIF2a adjusts survival and cell homeostasis in the ER stress mechanism (Pakos-zebrucka ve ark., 2016). Phosphorylated activation of EIF-α, which occurs in gestational diseases, especially preeclampsia, increases the SDF2 level of ER stress (Lorenzon-ojea, Wa, Burton, ve Bevilacqua, 2020).). It has been reported that maternal malperfusion causes ER stress in the placenta, resulting in intrauterine growth retardation, and acts with eIF2α phosphorylation through the AKT pathway (Yung ve ark., 2008).

As a result, it has been revealed that with the increase in HSPA5 gene expression and GRP78 protein level regarding the ER stress pathway, Thapsigargin affects intracellular calcium homeostasis in human trophoectoderm cells and creates ER stress in 2-dimensional cells and 3-dimensional spheroids. As the ER stress panel was interpreted with the findings obtained from the selected genes, it was seen that the ER stress induced by Thapsigargin in human trophectoderm cells was triggered by all 3 ER stress signaling pathways in 2- and 3-dimensional spheroid cells. In 2 dimensions, the process proceeds in the direction of EIF2A and ERN1, in 3 dimensions it proceeds in the direction of EIF2A and ATF6 and the increase in IRE1 protein level is evident in both experimental groups.

Misfolded proteins in the ER are directed to correct folding with the Unfolded Protein Response (UPR), primarily through GRP78, GRP94, calnexin, calreticulin and calcium-dependent chaperones. When misfolded proteins cannot be corrected, the amount of misfolded protein in the ER lumen increases and causes ER stress. As ER stress occurs, the cell tries to correct the misfolded proteins that cause stress by reducing the transcription of the misfolded proteins and inducing autophagy with the help of chaperones. Activated chaperones prompt different mechanisms and signal pathways and boost ways to relieve stress. Misfolded protein accumulation, which cannot be corrected even with activation of all these pathways, can conduct the cell to apoptosis (Xu, Bailly-Maitre, ve Reed, 2005). BCL2 gene codes a protein located in the outer mitochondrial membrane that blocks the apoptotic death of some cells (Czabotar, Lessene, Strasser, ve Adams, 2014). Ca^2+^ plays an important role in apoptosis induction linkage. It has been reported that the anti-apoptotic key Bcl-2 regulates the movement of Ca^2+^ across the ER membrane and its exact effect on calcium levels is controversial. Neverthless it is also stated that Ca^2+^ and Bcl-2 play a key role in induced apoptosis by ER stress. It has been reported that Bcl-2 regulates Ca^2+^ entry by SOCE (store-operated calcium entry) for ER stress-induced apoptosis modulation (Chiu ve ark., 2018). When Bcl2 expression levels were compared in trophoblasts of hyperglycemic and healthy placentas, it was shown that Bcl2 expression was decreased in trophoblast cells derived from hyperglycemic placenta, thus apoptosis was induced (Sgarbosa ve ark., 2006). It has been shown that in trophoblasts in recurrent pregnancy losses and preeclampsia all three pathways of ER stress are activated, apoptosis is induced with the decrease of Bcl2 and that endocannabinoid 2-arachidonoylglycerol affects ER stress and cell cycle in BeWo cell line and primary trophoblast cells (Teixeira ve Correia-da-silva, 2020). The protein coded by the CASP3 gene is a cysteine-aspartic acid protease that plays a central role in the execution stage of cell apoptosis. The encoded protein cleaves and inactivates the poly (ADP-ribose) polymerase while the sterol regulatory cleaves and activates the element binding proteins and caspases 6, 7 and 9. This protein itself is activated by Caspase 8, 9 and 10 (Porter ve Ja, 1999). Caspase 3 is mainly activated from caspases in ER stress and triggers apoptosis (Hitomi ve ark., 2004). As a result of malnutrition, ER stress is activated in trophoblast cells in the placenta. Also it has been shown that Caspase 3 is activated and apoptosis is induced in the cells (Arroyo ve ark., 2010). UPR response, IRE1, the UPR sensor, is activated via Caspas 3, and as a result, the expression of LDTM (ER-lumenal domain and transmembrane segment) increases. This expression is inhibited by apoptosis by inhibiting proapoptotic BAX in mitochondria (Shemorry, Harnoss, Guttman, ve Marsters, 2019). As a result of our analysis, while BCL2 expression showed a 16-fold increase in the 2-dimensional experimental group, no significant change was found in the 3-dimensional experimental group. In the CASP3 gene expression, upregulation occurred in the 2-dimensional experimental group compared to the control group, and downregulation occurred in the 3-dimensional spheroid experimental group compared to the control group. As a consequence, when the genes selected in the apoptosis panel and the findings obtained with IF staining are evaluated; Depending on the UPR response to ER stress induced by Thapsigargin, the 2 and 3 dimensional spheroid cells in the HTR-8/Vneo human trophoectoderm cell line show different characterizations. Cell survival mechanism with increased activation of apoptotic CASP3 and antiapoptotic BCL2 in 2 dimensional experimental group; significant increase in protein level of c-Caspase3 in experimental groups of both dimensions was evaluated as orientation to ER stress-mediated apoptotic cascade.

MAP1LC3B (Microtubule-associated proteins 1A / 1B light chain 3B) is a protein that in humans is encoded by the MAP1LC3B gene. LC3 is the most widely used marker of autophagosomes. LC3 is a central protein in the autophagy pathway where it functions in autophagy substrate selection and autophagosome biogenesis. It plays a role in mitophagy, which contributes to regulating mitochondrial quantity and quality by keeping mitochondria at basal level to meet cellular energy requirements and prevent ROS production. While LC3s are involved in the elongation of the phagophore membrane, the GABARAP/GATE-16 subfamily is involved in autophagosome maturation (Y. Lee ve Lee, 2016). It has been reported that UPR is regulated by autophagy signaling pathway in ER stress induced by the formation of UPR response (Kabir ve Kim, 2018) and apoptosis, autophagy and necrosis mechanisms are activated as a result of ER stress in villous trophoblasts collected from the placenta of preeclamptic pregnant women (Hutabarat, Wibowo, ve Huppertz, 2017). ER stress occurs in trophoblast cells isolated from patients with placental dysfunction and autophagy mechanisms due to lysosomal degradation in the cells are activated. In preeclampsia, ER stress caused by inflammatory cytokines released from trophoblast cells triggers the pyroptosis death pathway through autophagy mechanisms (S. Cheng ve ark., 2019). In our study, when MAP1LC3B was examined at the gene level, it was evaluated that there was no significant difference in the 2-dimensional experimental group and decreased by 2.2 times in the 3-dimensional experimental group. BECN1 gene codes a protein that regulates autophagy, the catabolic degradation process caused by intracellular starvation. The encoded protein vesicle is a component of the traffic-mediating phosphatidylinositol-3-kinase (PI3K) complex. This protein is thought to play a role in many cellular processes including tumorigenesis, neurodegeneration and apoptosis (Kang, Zeh, Lotze, ve Tang, 2011). Besides autophagy Beclin 1 plays a role in the proapoptotic process and Bcl2/Beclin1 interaction is important through the way that the cell will follow in the mechanisms of apoptosis and autophagy (Decuypere, Parys, ve Bultynck, 2012). Beclin 1 phosphorylation plays a major role in the autophagy mechanism (Menon ve Dhamija, 2018). It has been reported in the breast cancer cell line that autophagy mechanisms are activated to ensure cell homeostasis by increasing the protein level of LC3 and Beclin 1 in ER stress (X. I. U. Cheng, Liu, Jiang, Fang, ve Chen, 2014). In studies conducted in gestational trophoblast diseases, apoptosis and autophagy mechanisms were evaluated through Bcl-2 and Beclin1 immunoexpressions (Wargasetia, Shahib, Martaadisoebrata, Dhianawaty, ve Hernowo, 2015). ER stress occurs in the syncytialization processes of villous cytotrophoblasts and autophagy pathways are activated through Beclin 1 during the adaptation process (Bastida-ruiz ve ark., 2019). There was no significant change in BECN1 gene expression between the 2-dimensional experimental group and the control group. Upregulation was observed in the 3-dimensional experimental group compared to the control group. In Beclin1 IF analyzes associated with autophagy, the radiation intensity was found to be significantly higher in both groups at the protein level.

## 5. CONCLUSION

Here, when the ER stress panel was evaluated with the findings obtained from the selected genes, it was seen that all three ER stress signal pathways were triggered. The process in 2 dimensions progresses in the direction of EIF2A and ERN1, in 3 dimensions in the direction of EIF2A, ATF6 and the increase in IRE1 protein level in the two experimental groups is evident. As a result of ER stress, Beclin1-mediated autophagy and antiapoptotic Bcl2 are activated and survival mechanisms including the regulation of impaired calcium homeostasis are triggered. In addition, activation of c-Caspase3 at the protein level put forth ER stress-mediated apoptosis.

## 7. ACKNOWLEDGEMENTS

The authors would like to thank TUBITAK to financial support and Associate Professor Timur Kose for statistical analysis.

## 8. CONFLICT OF İNTEREST

The author declares that there is none of the conflicts.

## REFERENCES

Abou-Kheir, W., Barrak, J., Hadadeh, O., & Daoud, G. (2017). HTR-8/SVneo cell line contains a mixed population of cells. Placenta, 50(2017), 1–7. https://doi.org/10.1016/j.placenta.2016.12.007

Adams, C. J., Kopp, M. C., Larburu, N., Nowak, P. R., & Ali, M. M. U. (2019). Structure and Molecular Mechanism of ER Stress Signaling by the Unfolded Protein Response Signal Activator IRE1. Frontiers in Molecular Biosciences. Retrieved from https://www.frontiersin.org/article/10.3389/fmolb.2019.00011

Arroyo, J. A., Li, C., Schlabritz-Loutsevitch, N., McDonald, T., Nathanielsz, P., & Galan, H. L. (2010). Increased placental XIAP and caspase 3 is associated with increased placental apoptosis in a baboon model of maternal nutrient reduction. American Journal of Obstetrics and Gynecology, 203(4), 364.e13-364.e18. https://doi.org/10.1016/j.ajog.2010.05.021

Balahmar, R. M., Boocock, D. J., Coveney, C., Ray, S., Vadakekolathu, J., Regad, T., … Sivasubramaniam, S. (2018). Identification and characterisation of NANOG+/ OCT-4high/SOX2+ doxorubicin-resistant stem-like cells from transformed trophoblastic cell lines. Oncotarget, 9(6), 7054–7065. https://doi.org/10.18632/oncotarget.24151

Bastida-Ruiz, D., Yart, L., Wuillemin, C., Ribaux, P., Morris, N., & Epiney, M. (2019). The fine-tuning of endoplasmic reticulum stress response and autophagy activation during trophoblast syncytialization. Cell Death and Disease, 10(9). https://doi.org/10.1038/s41419-019-1905-6

Chen, Y., & Brandizzi, F. (2014). IRE1: ER stress sensor and cell fate executor. Trends Cell Biol. 2013 November, 23(11), 22–25. https://doi.org/10.1016/j.tcb.2013.06.005.IRE1

Cheng, S., Nakashima, A., Huber, W. J., Davis, S., Banerjee, S., & Huang, Z. (2019). Pyroptosis is a critical in fl ammatory pathway in the placenta from early onset preeclampsia and in human trophoblasts exposed to hypoxia and endoplasmic reticulum stressors. Cell Death and Disease. https://doi.org/10.1038/s41419-019-2162-4

Cheng, X. I. U., Liu, H. A. O., Jiang, C., Fang, L. I. N., & Chen, C. (2014). Connecting endoplasmic reticulum stress to autophagy through IRE1 / JNK / beclin-1 in breast cancer cells, 772–781. https://doi.org/10.3892/ijmm.2014.1822

Chiu, W., Chang, H., Lin, Y., Lin, Y., Chang, H., Lin, H., & Huang, S. (2018). Bcl-2 regulates store-operated Ca 2 + entry to modulate ER stress-induced apoptosis. Cell Death Discovery. https://doi.org/10.1038/s41420-018-0039-4

Collawn, J. F., & Bartoszewski, R. (2020). IRE1 Endoribonuclease Activity Modulates Hypoxic HIF-1 α Signaling in Human Endothelial Cells. Biomolecules, 10(6), 1–14.

Czabotar, P. E., Lessene, G., Strasser, A., & Adams, J. M. (2014). Control of apoptosis by the BCL-2 protein family: implications for physiology and therapy. Nature Reviews Molecular Cell Biology, 15(1), 49–63. https://doi.org/10.1038/nrm3722

De Paepe, C., Aberkane, A., Dewandre, D., Essahib, W., Sermon, K., Geens, M., … Van de Velde, H. (2018). BMP4 plays a role in apoptosis during human preimplantation development. Molecular Reproduction and Development. https://doi.org/10.1002/mrd.23081

Decuypere, J., Parys, J. B., & Bultynck, G. (2012). Regulation of the Autophagic Bcl-2/Beclin 1 Interaction, 284–312. https://doi.org/10.3390/cells1030284

Dicks, N., Bohrer, R. C., Gutierrez, K., Michalak, M., Agellon, L. B., & Bordignon, V. (2017). Relief of endoplasmic reticulum stress enhances DNA damage repair and improves development of pre-implantation embryos. PLOS ONE, 12(11), e0187717. Retrieved from https://doi.org/10.1371/journal.pone.0187717

Du, L., He, F., Kuang, L., Tang, W., Li, Y., & Chen, D. (2017). eNOS / iNOS and endoplasmic reticulum stress-induced apoptosis in the placentas of patients with preeclampsia, (October 2015), 49–55. https://doi.org/10.1038/jhh.2016.17

Elhussein, O. G., Ahmed, M. A., Suliman, S. O., Yahya, leena I., & Adam, I. (2019). Epidemiology of infertility and characteristics of infertile couples requesting assisted reproduction in a low-resource setting in Africa, Sudan. Fertility Research and Practice, 5(1), 7–11. https://doi.org/10.1186/s40738-019-0060-1

Gamage, T. K. J. B., Chamley, L. W., & James, J. L. (2016). Stem cell insights into human trophoblast lineage differentiation. Human Reproduction Update, 23(1), 77–103. https://doi.org/10.1093/humupd/dmw026

Ghosh, R., Lipson, K. L., Sargent, K. E., Mercurio, A. M., Hunt, J. S., & Urano, F. (2010). Transcriptional Regulation of VEGF-A by the Unfolded Protein Response Pathway, 5(3). https://doi.org/10.1371/journal.pone.0009575

Guzel, E., Arlier, S., Guzeloglu-Kayisli, O., Tabak, M. S., Ekiz, T., Semerci, N., … Kayisli, U. A. (2017). Endoplasmic reticulum stress and homeostasis in reproductive physiology and pathology. International Journal of Molecular Sciences, 18(4). https://doi.org/10.3390/ijms18040792

Hannan, N. J., Paiva, P., Dimitriadis, E., & Salamonsen, L. A. (2010). Models for Study of Human Embryo Implantation: Choice of Cell Lines?1. Biology of Reproduction, 82(2), 235–245. https://doi.org/10.1095/biolreprod.109.077800

He, B., Zhang, H., Wang, J., Liu, M., Sun, Y., Guo, C., … Kong, S. (2019). Blastocyst activation engenders transcriptome reprogram affecting X-chromosome reactivation and inflammatory trigger of implantation. Proceedings of the National Academy of Sciences, 116(33), 16621 LP –16630. https://doi.org/10.1073/pnas.1900401116

Hebert-schuster, M., Rotta, B. E., Kirkpatrick, B., Guibourdenche, J., & Cohen, M. (2018). The Interplay between Glucose-Regulated Protein 78 (GRP78) and Steroids in the Reproductive System, 78, 1–11. https://doi.org/10.3390/ijms19071842

Hillary, R. F., & Fitzgerald, U. (2018). A lifetime of stress : ATF6 in development and homeostasis, 1–10.

Hitomi, J., Katayama, T., Taniguchi, M., Honda, A., Imaizumi, K., & Tohyama, M. (2004). Apoptosis induced by endoplasmic reticulum stress depends on activation of caspase-3 via caspase-12. Neuroscience Letters, 357(2), 127–130. https://doi.org/10.1016/j.neulet.2003.12.080

Hutabarat, M., Wibowo, N., & Huppertz, B. (2017). The trophoblast survival capacity in preeclampsia, 1–21.

Iurlaro, R., & Muñoz-Pinedo, C. (2016). Cell death induced by endoplasmic reticulum stress. FEBS Journal, 283, 2640–2652. https://doi.org/10.1111/febs.13598

Iwawaki, T., Akai, R., Yamanaka, S., & Kohno, K. (2009). Function of IRE1 alpha in the placenta is essential for, 106(39), 16657–16662.

Kabir, M. F., & Kim, H. (2018). Endoplasmic Reticulum Stress and Autophagy. IntechOpen, 1–24.

Kang, R., Zeh, H. J., Lotze, M. T., & Tang, D. (2011). The Beclin 1 network regulates autophagy and apoptosis. Cell Death and Differentiation, 18(4), 571–580. https://doi.org/10.1038/cdd.2010.191

Kim, S.-M., & Kim, J.-S. (2017). A Review of Mechanisms of Implantation. Development & Reproduction, 21(4), 351–359. https://doi.org/10.12717/dr.2017.21.4.351

Lee, C.-L., Veerbeek, J. H. W., Rana, T. K., van Rijn, B. B., Burton, G. J., & Yung, H. W. (2019). Role of Endoplasmic Reticulum Stress in Proinflammatory Cytokine–Mediated Inhibition of Trophoblast Invasion in Placenta-Related Complications of Pregnancy. The American Journal of Pathology, 189(2), 467–478. https://doi.org/10.1016/j.ajpath.2018.10.015

Lee, Y., & Lee, J. (2016). Role of the mammalian ATG8 / LC3 family in autophagy : differential and compensatory roles in the spatiotemporal regulation of autophagy, 49(8), 424–430.

Lorenzon-ojea, A. R., Wa, H., Burton, G. J., & Bevilacqua, E. (2020). BBA - Molecular Basis of Disease The potential contribution of stromal cell-derived factor 2 (SDF2) in endoplasmic reticulum stress response in severe preeclampsia and labor-. BBA - Molecular Basis of Disease, 1866(2), 165386. https://doi.org/10.1016/j.bbadis.2019.01.012

Malhotra, J. D., & Kaufman, R. J. (2007). The endoplasmic reticulum and the unfolded protein response. Seminars in Cell & Developmental Biology, 18(6), 716–731. https://doi.org/10.1016/j.semcdb.2007.09.003

Menon, M. B., & Dhamija, S. (2018). Beclin 1 Phosphorylation –at the Center of Autophagy Regulation. Frontiers in Cell and Developmental Biology, 6, 137. https://doi.org/10.3389/fcell.2018.00137

Michalak, M., & Gye, M. C. (2015). Endoplasmic reticulum stress in periimplantation embryos. Clinical and Experimental Reproductive Medicine, 42(1), 1–7. https://doi.org/10.5653/cerm.2015.42.1.1

Mochan, S., Dhingra, M. K., Varghese, B., & Gupta, S. K. (2017). sFlt-1 (sVEGFR1) induces placental endoplasmic reticulum stress in trophoblast cell : implications for the complications in preeclampsia-an in vitro study, 1.

Msheik, H., Azar, J., El Sabeh, M., Abou-Kheir, W., & Daoud, G. (2020). HTR-8/SVneo: A model for epithelial to mesenchymal transition in the human placenta. Placenta, 90, 90–97. https://doi.org/10.1016/j.placenta.2019.12.013

Nakashima, A., Cheng, S., Kusabiraki, T., Motomura, K., & Aoki, A. (2019). Endoplasmic reticulum stress disrupts lysosomal homeostasis and induces blockade of autophagic flux in human trophoblasts. Scientific Reports, (January), 1–12. https://doi.org/10.1038/s41598-019-47607-5

Nandi, P., Lim, H., Torres-Garcia, E. J., & Lala, P. K. (2018). Human trophoblast stem cell self-renewal and differentiation: Role of decorin. Scientific Reports, 8(1), 1–14. https://doi.org/10.1038/s41598-018-27119-4

Ozcan, L., & Tabas, I. (2012). Role of Endoplasmic Reticulum Stress in Metabolic Disease and Other Disorders. Annual Review of Medicine, 63(1), 317–328. https://doi.org/10.1146/annurev-med-043010-144749

Pakos-zebrucka, K., Koryga, I., Mnich, K., Ljujic, M., Samali, A., Gorman, A. M., & Pakos-zebrucka, K. (2016). The integrated stress response, 17(10), 1374–1395.

Porter, A. G., & Ja, R. U. (1999). Emerging roles of caspase-3 in apoptosis, 99–104.

Sgarbosa, F., Fernando, L., Brasil, M. A. M., Gonc, R., Costa, E., & Calderon, I. M. P. (2006). Changes in apoptosis and Bcl-2 expression in human hyperglycemic, term placental trophoblast, 1–7. https://doi.org/10.1016/j.diabres.2005.12.014

Shemorry, A., Harnoss, J. M., Guttman, O., & Marsters, S. A. (2019). Caspase-mediated cleavage of IRE1 controls apoptotic cell commitment during endoplasmic reticulum stress. ELife 2019;8:E47084, 1, 1–23.

Song, W., Chang, W.-L., Shan, D., Gu, Y., Gao, L., Liang, S., … Liu, X. (2020). Intermittent Hypoxia Impairs Trophoblast Cell Viability by Triggering the Endoplasmic Reticulum Stress Pathway. Reproductive Sciences, 27(2), 477–487. https://doi.org/10.1007/s43032-019-00039-y

Takao, T., Asanoma, K., Kato, K., Fukushima, K., Tsunematsu, R., Hirakawa, T., … Wake, N. (2011). Isolation and characterization of human trophoblast side-population (SP) cells in primary villous Cytotrophoblasts and HTR-8/SVneo cell line. PLoS ONE, 6(7). https://doi.org/10.1371/journal.pone.0021990

Tatar, M., & Tatar, T. (2018). Endoplazmik Retikulum Stresi ve İlişkili Hastalıklar. OSMANGAZİ Journal of Medicine, 00, 294–303. https://doi.org/10.20515/otd.417682

Teixeira, N., & Correia-da-silva, G. (2020). The endocannabinoid 2-arachidonoylglycerol promotes endoplasmic reticulum stress in placental cells.

Wargasetia, T. L., Shahib, M. N., Martaadisoebrata, D., Dhianawaty, D., & Hernowo, B. (2015). Characterization of apoptosis and autophagy through Bcl-2 and Beclin-1 immunoexpression in gestational trophoblastic disease, 13(7), 413–420.

Weber, M., Knoefler, I., Schleussner, E., Markert, U. R., & Fitzgerald, J. S. (2013). HTR8/SVneo cells display trophoblast progenitor cell-like characteristics indicative of self-renewal, repopulation activity, and expression of “stemness-” associated transcription factors. BioMed Research International, 2013. https://doi.org/10.1155/2013/243649

Winship, A., Ton, A., Van Sinderen, M., Menkhorst, E., Rainczuk, K., Griffiths, M., … Dimitriadis, E. (2018). Mouse double minute homologue 2 (MDM2) downregulation by miR-661 impairs human endometrial epithelial cell adhesive capacity. Reproduction, Fertility and Development, 30(3), 477–486. https://doi.org/10.1071/RD17095

Xu, C., Bailly-Maitre, B., & Reed, J. (2005). Endoplasmic reticulum stress: cell life and death decisions. Journal of Clinical Investigation, 115(10), 2656–2664. https://doi.org/10.1172/JCI26373.2656

Yang, C., Lim, W., Bazer, F. W., & Song, G. (2017). Propyl gallate induces cell death and inhibits invasion of human trophoblasts by blocking the AKT and mitogen-activated protein kinase pathways. Food and Chemical Toxicology, 109 (August), 497–504. https://doi.org/10.1016/j.fct.2017.09.049

Yang, C., Lim, W., Bazer, F. W., & Song, G. (2018). Butyl paraben promotes apoptosis in human trophoblast cells through increased oxidative stress-induced endoplasmic reticulum stress. Environmental Toxicology, 33(4), 436–445. https://doi.org/10.1002/tox.22529

Yung, H., Calabrese, S., Hynx, D., Hemmings, B. A., Cetin, I., Charnock-jones, D. S., & Burton, G. J. (2008). Evidence of Placental Translation Inhibition and Endoplasmic Reticulum Stress in the Etiology of Human Intrauterine Growth Restriction, 173(2), 451–462. https://doi.org/10.2353/ajpath.2008.071193

